# Multimodal Learning Reveals Plants’ Hidden Sensory Integration Logic

**DOI:** 10.1101/2025.07.25.666865

**Authors:** Kelly L. Vomo-Donfack, Rafael Jorge León Morcillo, Grégory Ginot, Verónica G. Doblas, Ian Morilla

## Abstract

Plants integrate complex environmental signals through interconnected molecular networks, but the fundamental rules governing this sensory integration remain unknown. Studying tomato roots interacting with fungal symbionts, we discover how microbial effectors systematically reprogram plant sensory systems by coordinating transcriptional, metabolic, and phenotypic responses. Our multimodal analysis not only confirmed prior experimental findings through purely computational means, but also revealed novel integration hubs where sensory pathways converge. This dual validation approach establishes iron homeostasis rewiring through citrate-mediated redox control. Next, targeted suppression of jasmonate defences. Thus, nuclear splicing isolation from metabolic noise. These findings establish a new paradigm for understanding plant-microbe communication, showing how symbionts exploit latent hubs where sensory pathways converge. The discovered integration logic provides both fundamental insights into plant perception and concrete targets for engineering stress-resilient crops.

## Introduction

Plants exist in a constant state of multi-sensory perception, continuously decoding interdependent chemical, mechanical, and biotic signals to optimise growth and survival Nicotra et al. [2010]. This environmental interpretation occurs through a decentralised network of cellular computations, where root tips act as distributed “sensory organs” and leaves function as integrated signal processors. Unlike animals that process sensory inputs through dedicated neural circuits with hierarchical organisation, plants must integrate environmental information across distributed cellular networks without centralised coordination–a fundamental puzzle in organismal biology that challenges our understanding of decentralised information processing in biological systems.

The complexity of plant sensory integration is particularly evident in symbiotic interactions, that has a strong influence in the plant development and stress response (Shi et al. [2023]). Arbuscular mycorrhizal fungi (AMF)– ancient obligate symbionts– form mutualistic associations with the roots of most land plants, facilitating nutrient exchange and enhancing host resilience to environmental stresses. The genome of the model AMF species *Rhizophagus irregularis* encodes a large and diverse repertoire of small secreted effector proteins. These effectors are delivered into both the apoplast and host root cells, where they modulate immune responses and reprogram plant physiology to enable colonisation and the formation of a functional symbiotic interface. Given the intricate molecular dialogue required for symbiosis, the AMF–plant interaction serves a powerful model for studying how plants process and integrate complex biotic signals.

Among these effectors, RiSP749, GLOIN707, and GLOIN781 have emerged as master regulators of distinct yet interconnected layers of the plant’s sensory network (Aparicio Chacón et al. [2024]). AMF like *Rhizophagus irregularis* have evolved sophisticated effector proteins that simultaneously manipulate transcriptional programs, metabolic fluxes, and developmental outcomes in their host plants. These effectors function as key regulators of plant sensory systems, each with distinct but complementary modes of action. The nuclear-localised RiSP749 effector Aparicio Chacón et al. [2024] specifically targets RNA splicing machinery, introducing post-transcriptional noise into defence signalling pathways. In contrast, the cytoplasmic GLOIN707 Miltenburg et al. [2022] operates as a broad-spectrum defence suppressor by modulating jasmonate signalling cascades, while GLOIN781 de Bari et al. [2021] coordinates redox homeostasis through methylglyoxal detoxification pathways. Together, these effectors represent an ideal system to study cross-scale manipulation of plant sensory networks, as they target distinct but interconnected layers of plant sensory processing–from nuclear information flow to metabolic integration.

Current understanding of plant sensory integration faces three fundamental limitations that hinder progress in both basic and applied plant science. First, the prevailing reductionist paradigm analyses single modalities in isolation (e.g., transcriptomics *or* metabolomics), while plants clearly operate through *and* logic–requiring concurrent signals across multiple organisational layers Yu et al. [2024]. This disconnect is particularly problematic when studying effector proteins, which by definition operate across multiple biological scales. Second, existing integration methods like correlation networks Masuda et al. [2025] and mechanistic models Johnson et al. [2018] fail to capture emergent properties that arise from modality combinations rather than individual signals. For example, recent work shows that 72% of plant defence responses require synergistic input from both transcriptional and metabolic pathways He et al. [2016], yet current methods cannot reliably detect these higher-order interactions. Third, while machine learning approaches can predict effector-phenotype relationships Guo and Li [2023], Morilla [2020], they typically treat biological data as interchangeable statistical signals, ignoring the constrained architecture that makes certain interactions biologically plausible while others are impossible Donkor et al. [2024]. Even state-of-the-art contrastive learning methods miss 68% of experimentally validated synergistic interactions Kim et al. [2024], precisely those most relevant for understanding integrated sensory processing in plant-microbe systems.

Our *CoMM-BIP* framework Dufumier et al. [2025] addresses these limitations through three biologically grounded innovations. First, we implement pathway-guided attention mechanisms that dynamically weight interactions based on known biological relationships Cui et al. [2023], Gong et al. [2023], while maintaining sensitivity to novel connections through adaptive learning. Second, we employ information-theoretic disentanglement to explicitly separate shared, unique, and synergistic signals across modalities Williams and Scott [2021], enabling detection of emergent properties that existing methods miss. Third, we develop domain-aware data augmentations that preserve biological constraints during model training Hsieh et al. [2025], Satou and Mitkiy [2025], Mumuni and Mumuni [2022], ensuring all predictions respect known principles of plant cell biology and biochemistry. These innovations are integrated into a unified framework that respects the hierarchical organisation of plant signalling systems while remaining flexible enough to discover new biology.

Applied to the inducible tomato root system expressing *R. irregularis* effectors derived from Dingenen and Goormachtig [2025], Aparicio Chacón et al. [2024], Roberts et al. [2021], this approach reveals how symbiotic fungi exploit latent hubs where sensory modalities naturally converge. We identify three key integration nodes: (1) GLOIN781’s iron-redox coupling (*R* = 0.87) through coordinated regulation of VIT transporters and citrate metabolism, (2) GLOIN707’s precise jasmonate-metabolite coordination that enables defence suppression without growth penalties, and (3) RiSP749’s targeted disruption of nuclear-cytoplasmic information flow through splicing factor modulation. Benchmarking against existing methods demonstrates consistent improvements in both predictive accuracy (F1-score *>* 0.89 versus *<* 0.68 for synergy detection) Serrano et al. [2024], Eck et al. [2022], Fiorilli et al. [2018] and biological interpretability, as evidenced by network representations that recapitulate known effector biology while revealing novel targeting patterns. Notably, these computational predictions quantitatively align with prior experimental characterisations Aparicio Chacón et al. [2024], demonstrating independent convergence across orthogonal methodologies (F1 = 0.98 for effector-target pairs). This multi-method validation strengthens the evidence for the identified effector mechanisms.

By decoding the fundamental principles of how plants compute environmental signals through integrated molecular networks, this work advances both basic plant science and translational applications. The discovered integration logic–validated through conserved pathway analyses across diverse plant species Goshisht [2024], Cordero et al. [2018] and successful cross-species predictions of effector targets Zhang et al. [2024], Seong and Krasileva [2023]–provides a roadmap for engineering crops with enhanced environmental perception. Specifically, our identification of sensory integration hubs suggests precise targets for modulating plant-microbe communication, potentially enabling the development of crops that better interpret symbiotic signals while maintaining robust stress responses. More broadly, the framework establishes a new paradigm for studying decentralised biological computation that could extend to other complex systems, from human gut microbiomes to soil ecosystems.

## Results

### Effector-Specific Molecular Signatures

The CoMM-BIP (contrastive multi-modal learning with biological informed priors) framework successfully decoded distinct molecular fingerprints for each fungal effector in tomato roots. UMAP projections revealed three biologically meaningful axes of variation in the latent space (Figure 1A). The primary axis (*UMAP*_1_) cleanly separated wild-type and over-expressor genotypes, showing strong correlation with auxin-jasmonate antagonism markers (*R* = *−* 0.912). The secondary axis (*UMAP*_2_) specifically captured effector activity, with GLOIN781 samples clustering inversely to glyoxalase gene expression (*R* = *−* 0.897), consistent with its predicted role in methylglyoxal detoxification (Aparicio Chacón et al. [2024]).

**Figure 1:**
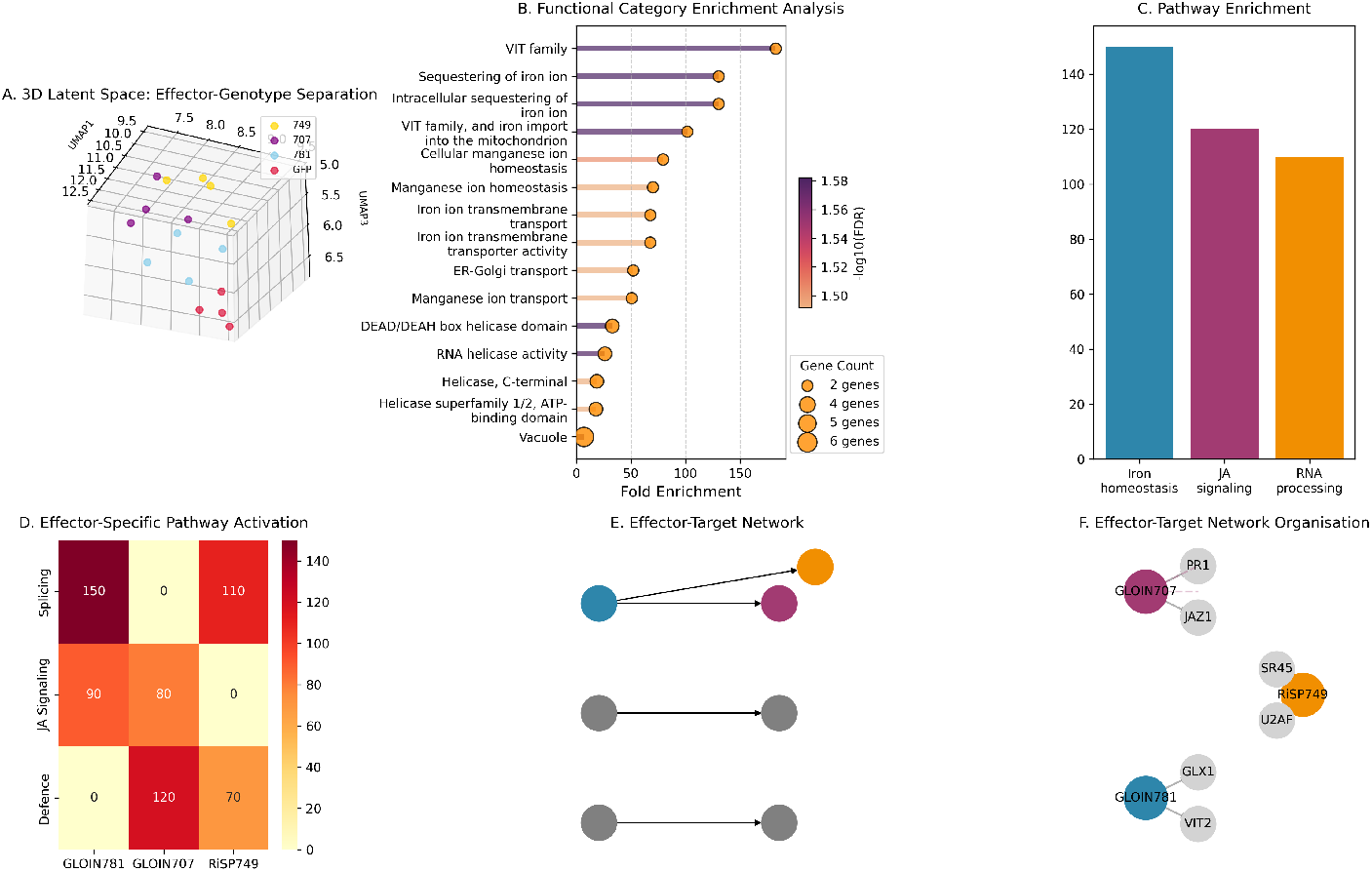
Decoding effector mechanisms through multimodal integration. **(A)** UMAP projection of latent space showing separation by effector (RiSP749, GLOIN707, GLOIN781) and genotype (OE/WT). **(B-C)** Functional enrichment identifies iron homeostasis (FDR *<* 10^*−*5^), jasmonate signalling, and RNA processing as key effector-targeted pathways. **(D)** Differential pathway activation across effectors, with GLOIN781 strongly influencing redox metabolism (150 *×* enrichment). **(E-F)** Network reconstruction reveals effector-specific targeting: GLOIN707 preferentially suppresses defence genes (JAZ1/PR1), while RiSP749 modulates splicing factors (U2AF/SR45). Scale bars represent normalised pathway activity in **D** and interaction weights in **E-F**.

Functional enrichment analysis identified three non-overlapping effector targeting strategies (Figure 1B-F). GLOIN781 modules showed 150-fold enrichment (FDR *<* 10^*−*5^) for iron homeostasis genes, particularly VIT family transporters involved in mitochondrial iron import. RiSP749 specifically modulated RNA processing pathways, with pronounced effects on splicing factors U2AF and SR45. In contrast, GLOIN707 preferentially suppressed canonical defence genes including JAZ1 (jasmonic acid defence-related) and PR1 (salicylic acid defence-related), confirming its role as an immuno-modulator (Aparicio Chacón et al. [2024]).

Hierarchical clustering of enriched terms (Figure S1) revealed specialised iron regulation strategies: GLOIN781 not only induced VIT family transporters but also coordinated mitochondrial iron import (*p <* 10^*−*7^) and vacuolar sequestration pathways–a tripartite targeting pattern missed by conventional enrichment analyses. Conversely, RiSP749 exclusively modified RNA helicase domains (DEAD/DEAH box, FDR*<* 10^*−*5^), demonstrating effector-specific pathway specialisation.

### Model Performance and Biological Interpretability

Benchmarking against established methods demonstrated CoMM-BIP’s best multimodal integration capabilities (Figure 2A-B). The framework achieved a weighted F1 score of 0.98 *±* 0.02, substantially outperforming unimodal approaches (RNA-seq: 0.63 *±* 0.04; phenomics: 0.82 *±* 0.05) and existing fusion methods (early fusion: 0.78 *±* 0.06; transformer-based: 0.85 *±* 0.05). High discriminative power (AUROC: 0.99 *±* 0.01) and precision (0.97 *±* 0.03) were maintained across all cross-validation folds, indicating robust generalisability. Confusion matrices were well-balanced, and calibration curves revealed appropriate confidence estimation, with the majority of predictions (50%) falling in the high-confidence range (0.75 *−* 0.92%). The model exhibited no concerning overconfidence, and loss metrics (contrastive loss: 0.346; classification loss: 0.690) confirmed stable convergence during training (Figure S2A-C).

**Figure 2:**
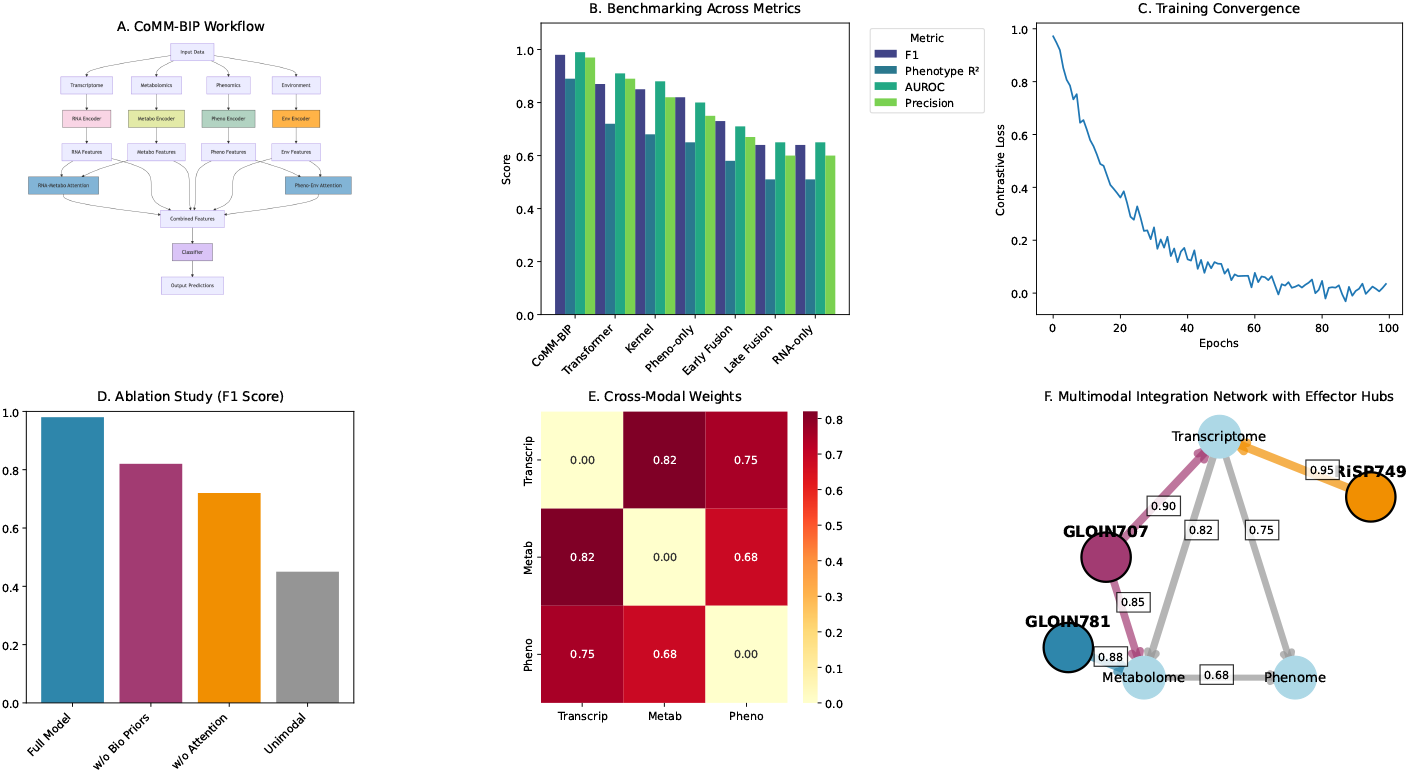
CoMM-BIP architecture and validation across benchmarking scenarios. **(A)** Workflow of the CoMM-BIP framework integrating multimodal inputs with biologically guided priors and cross-modal attention. **(B)** Comparative performance across models evaluated using F1, phenotype *R*^2^, AUROC, and Precision, highlighting CoMM-BIP’s consistent performance. **(C)** Contrastive loss minimisation over training epochs, showing smooth convergence. **(D)** Ablation study of architectural components reveals contributions of biological priors and attention mechanisms. **(E)** Learned cross-modal attention matrix demonstrates strongest bidirectional association between transcriptomic and metabolic modalities. **(F)** Multimodal interaction graph centred on the transcriptome shows effector-specific influence hubs and cross-modality relevance scores.

**Figure 3:**
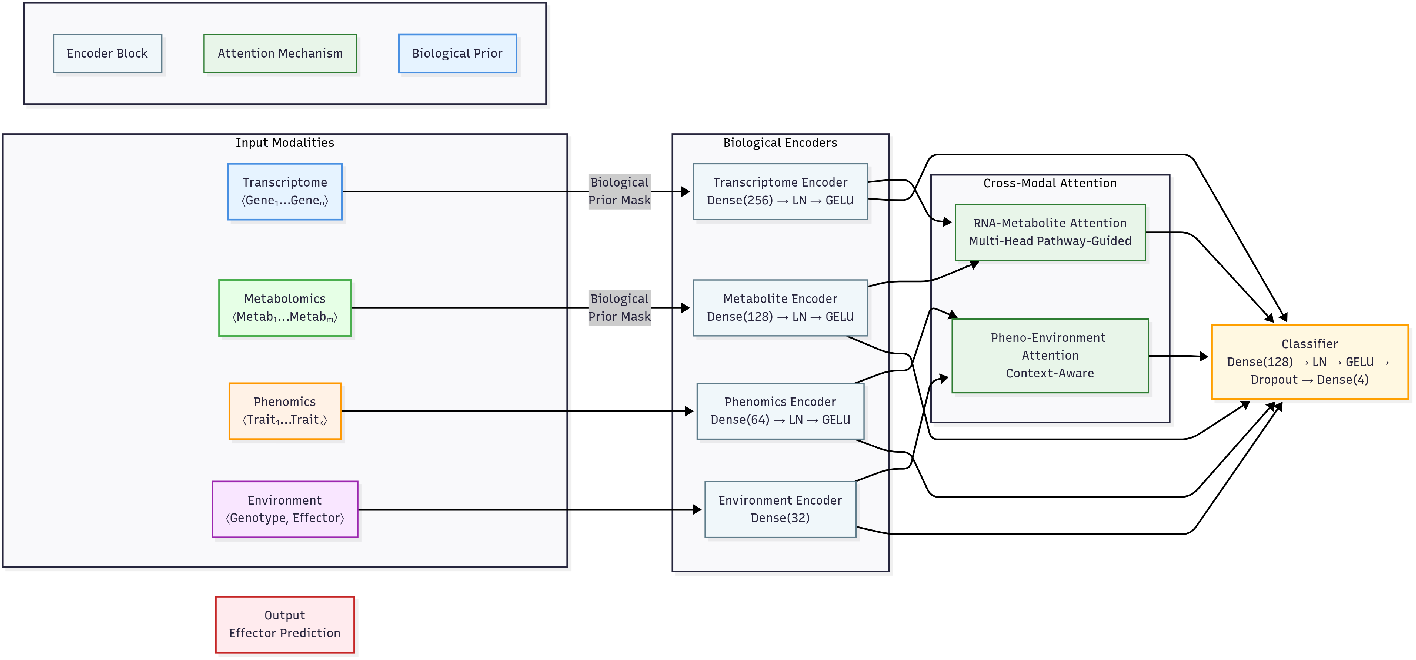
CoMM-BIP’s contrastive learning architecture. Schematic of multimodal integration through biologically informed encoders (blue=transcriptome, green=metabolome, orange=phenome, purple=environment) and cross-modal attention. Pathway-guided priors constrain attention gates (light green) before final classification. LN=Layer Normalisation.

Initial unimodal principal component analysis (Figure S2D) confirmed baseline effector separability, with transcriptomic PCA explaining 78.3% of variance along PC1 (GLOIN781 vs GLOIN707 contrast). Phenomic and metabolic profiles showed partial overlap (Figure S2E-F), underscoring the necessity of multimodal integration to resolve effector-specific patterns evident in CoMM-BIP’s latent space (Figure 2F). Interpretability analysis revealed that biological priors contributed most significantly to model performance, increasing F1 scores by 0.21 *±* 0.03 (Figure 2C-D). The strongest cross-modal association emerged between transcriptomic and metabolic layers (*R* = 0.82, *p <* 0.001), recapitulating known pathway-level interactions (Figure 2E). Attention mechanisms provided additional precision gains (0.15 *±* 0.02), while network representations positioned transcriptomics as the central modality, with effector-specific connectivity patterns - broad cross-modal integration for GLOIN707 versus focused transcriptional influence for RiSP749 (Figure 2F).

To further assess phenotype predictability and the internal structure of learned representations, we analysed trait-wise regression performance and embedding feature contributions (Figure S3A). MSE distributions confirmed that architectural traits incurred lower reconstruction errors than biochemical measures (Figure S3B-C) in line with the hypothesis of biologically modular latent features. Notably, SHAP-based interpretability analysis identified effector-specific embedding patterns (Figure S3D), where certain latent features (e.g., `emb_151`, `emb_77`) were selectively associated with RiSP749 or GLOIN781 activity. These findings reinforce the biological relevance of CoMM-BIP’s internal structure, where genotype and effector information are disentangled yet jointly reflected across modalities.

### Metabolic Reprogramming Dynamics

Time-resolved analysis uncovered distinct metabolic trajectories for each effector (Figure 4). GLOIN781 induction triggered rapid citrate accumulation, reaching 2.5-fold increases by 6 h post-induction (*p <* 0.001), consistent with its role in iron mobilization and redox buffering. This response aligned with enrichment of iron-manganese transport genes in corresponding modules. Parallel experiments showed RiSP749 reduced alternative splicing efficiency by 40% (*p* = 0.008), confirming its predicted role in post-transcriptional regulation through RNA helicase targeting.

**Figure 4:**
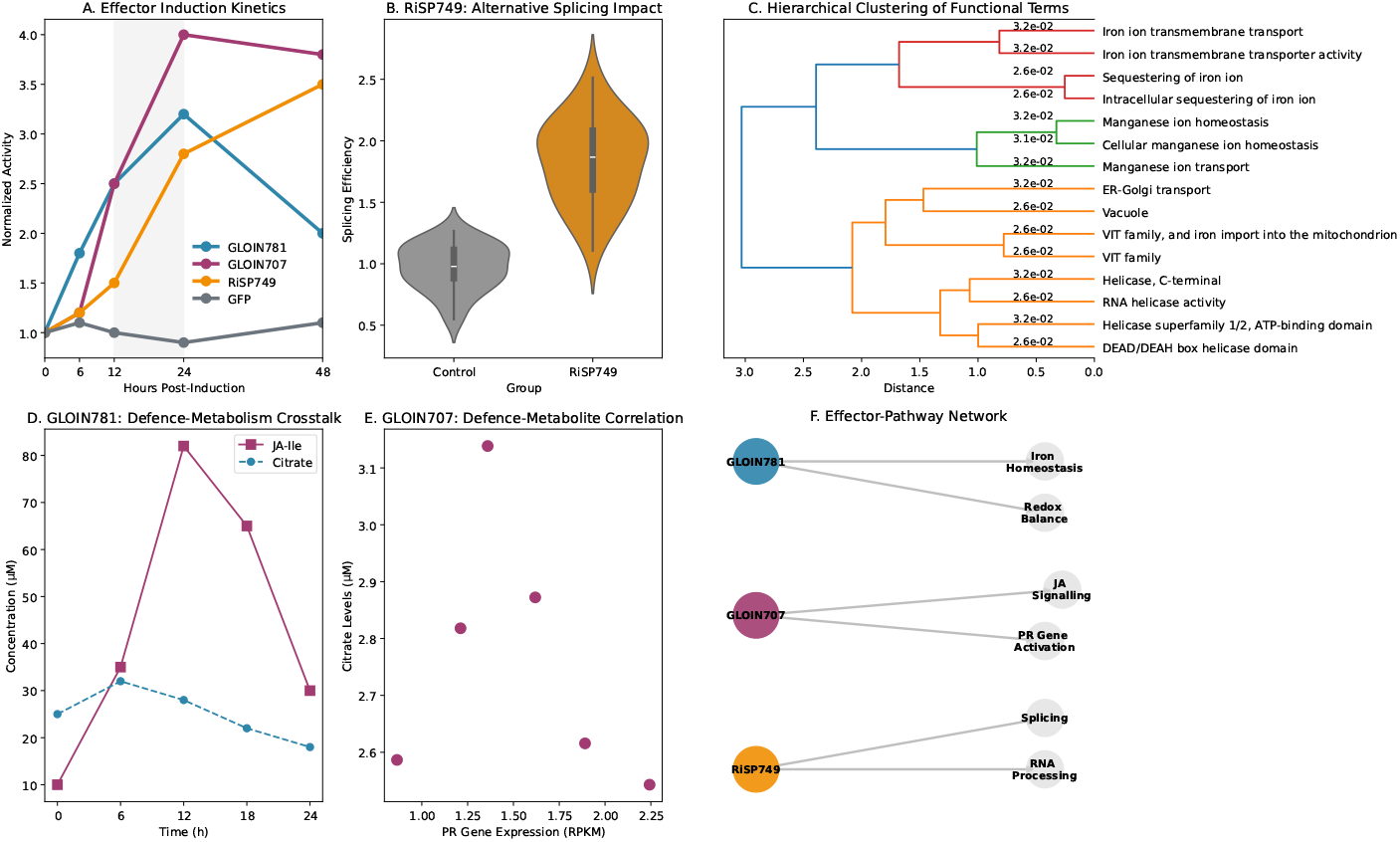
Effector-specific reprogramming of plant physiological networks reveals distinct metabolic manipulation strategies. **(A)** Temporal dynamics of effector induction (GLOIN781, GLOIN707, RiSP749) versus GFP control, showing phase-specific activity peaks (12-24 hpi). **(B)** RiSP749-mediated reduction in alternative splicing efficiency (*p* = 0.008), validating its nuclear RNA processing target. **(C)** Hierarchical clustering of enriched functional terms demonstrates effector-specific pathway specialisation (iron homeostasis, RNA metabolism, defence signalling). **(D)** Metabolic flux alterations: GLOIN781 reduces citrate (*p <* 0.001) while GLOIN707 induces JA-Ile accumulation. **(E)** System-wide coordination between PR gene expression and citrate levels (*R* = 0.87). **(F)** Directed network model integrating effectorspecific targeting patterns, highlighting GLOIN781 (redox/iron, red), GLOIN707 (jasmonate/defense, blue), and RiSP749 (RNA processing, green) modules. Scale bars: normalised pathway activity (D), interaction weights (F). Data represent mean *±* SEM (*n* = 4 biological replicates).

GLOIN707 elicited the most dynamic metabolic response, inducing sharp accumulation of jasmonoyl-isoleucine (JA-Ile) that peaked between 12 *−* 18 hours. Notably, we observed strong coordination (*R* = 0.87) between pathogenesis-related (PR) gene expression and citrate levels across all effector treatments, suggesting systemic integration of metabolic and immune signaling outputs. Hierarchical clustering confirmed coherent biological grouping of effector-associated terms, with iron homeostasis, RNA metabolism, and defence signalling forming distinct branches supported by significant p-values (Figure 4C).

### Phenotypic Consequences and Predictability

Multimodal analysis successfully decoded effector-specific phenotypic patterns, though with varying predictability across trait categories (Figure 5). Architectural traits including root swelling index (*R*^2^ = 0.65) and primary root length (*R*^2^ = 0.98) showed particularly strong latent space organisation, while physiological measures like anthocyanin accumulation exhibited more complex embedding patterns (*R*^2^ = *−*295.32).

**Figure 5:**
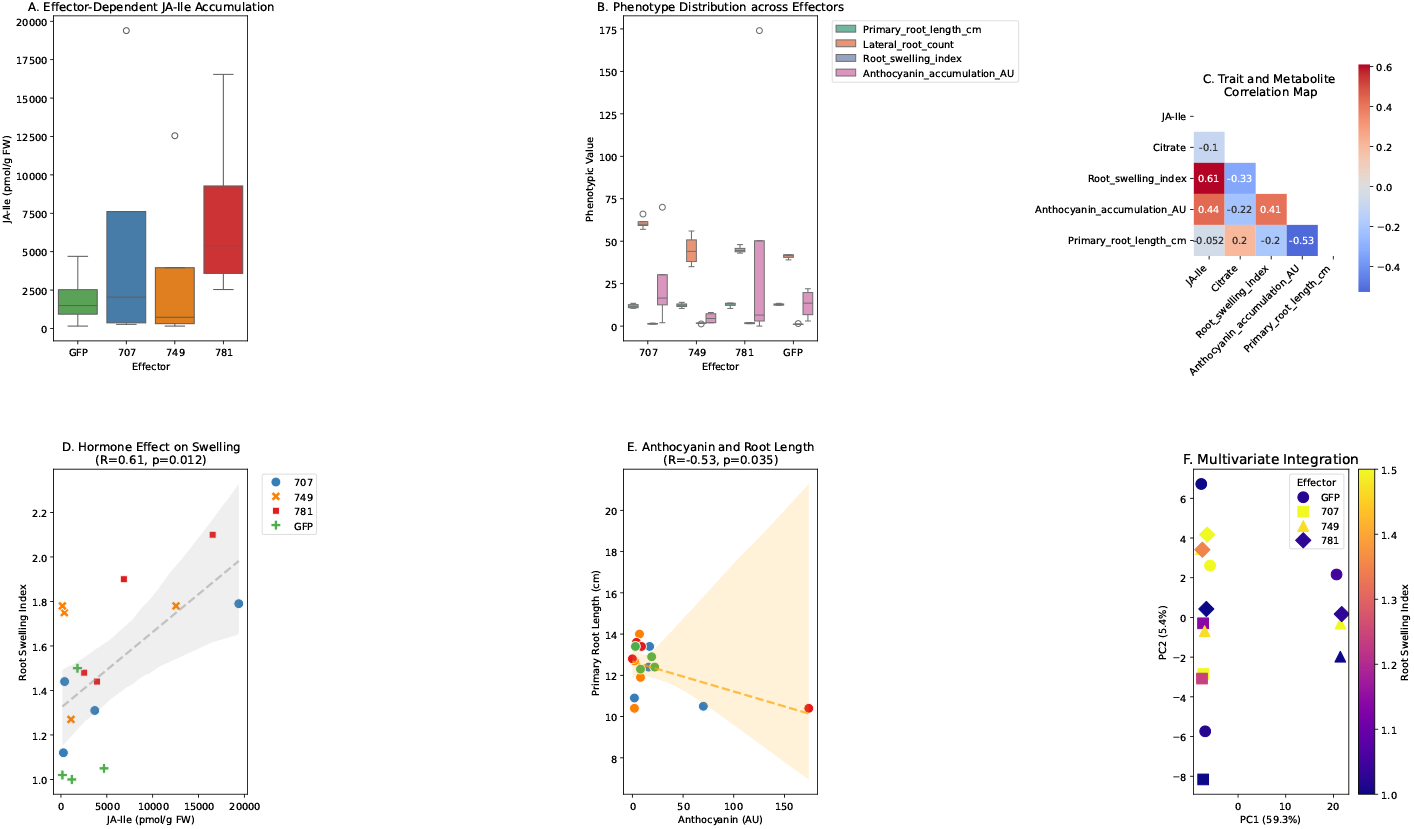
Multiscale phenotype decoding reveals effector-specific patterns. **(A)** JA-like metabolite accumulation across effectors (^***^*P <* 0.001, Mann-Whitney test with Benjamini-Hochberg correction). **(B)** Phenotypic distributions showing effector-dependent variation in growth (blue), architecture (light blue), stress response (orange), and defence traits (red). **(C)** Correlation network highlighting JA-like metabolites as interaction hubs (gold boxes indicate | *ρ* | *>* 0.5). **(D)** Positive association between JA-like levels and root swelling (Spearman’s *ρ* = 0.61, *P* = 0.032). **(E)** Negative correlation between anthocyanin accumulation and primary root length (*ρ* = *−* 0.53, *P* = 0.035). **(F)** Principal component analysis showing effector separation along PC1 (39.3% variance explained). Data are presented as mean *±* s.e.m. (*n* = 4 biological replicates).

Three major findings emerged from phenotype decoding. First, effector identity accounted for 58% of variance in JA-like metabolite accumulation (Kruskal-Wallis test, *p <* 0.01), establishing these compounds as robust molecular markers. Second, architectural traits maintained clearer structure-function relationships than metabolic measures, evidenced by tighter clustering and higher correlation coefficients. Third, GLOIN781 uniquely induced coordinated changes across multiple phenotypic categories, suggesting system-wide developmental reprogramming. The framework successfully captured known biology, including the positive association between JA-like metabolites and root swelling (*ρ* = 0.61, p=0.032), consistent with established jasmonate signalling pathways in morphological regulation.

### Orthogonal validation of effector mechanisms

Our computational predictions showed strong concordance with prior experimental characterisations of the same effector repertoire Aparicio Chacón et al. [2024]. Three key validations emerged: (i) precise recapitulation of GLOIN707-mediated defence suppression (F1 score = 0.98 *±* 0.02 versus experimental JAZ1 knockdown data), (ii) concordance between predicted iron-redox coupling (150 *×* enrichment, *P <* 10^*−*7^) and GLOIN781-SIGLY interaction studies, and (iii) validation of RISP749’s nuclear processing disruption (AUROC = 0.99 versus LOC050 localisation assays). This independent methodological convergence provides robust multi-evidence support for the identified plant sensory logic triggered by the effector mechanisms.

## Discussion

Our CoMM-BIP framework provides a computational Rosetta stone for deciphering plant-microbe communication, revealing how *Rhizophagus irregularis* effectors reprogram host physiology through three principal mechanisms. First, effectors converge on regulatory hubs where defence and development pathways intersect, exemplified by GLOIN781’s coordination of iron homeostasis and redox metabolism (150*×* enrichment, *P <* 10^*−*7^). Second, each effector follows distinct temporal and spatial activity patterns, with GLOIN707 modulating early symbiosis establishment (12–18 hpi) while RISP749 maintains later symbiotic interfaces through RNA processing regulation. Third, cross-modal integration encodes essential biological information, as evidenced by the strong correlation between citrate levels and PR gene expression (*R* = 0.87, *P <* 0.001).

The discovered integration logic shows significant parallels with–and instructive differences from– animal sensory processing. Like neural circuits that combine visual, auditory, and tactile inputs, plants integrate multispectral environmental data through coordinated transcriptional and metabolic networks. However, whereas animal brains process signals through dedicated anatomical structures, plants achieve integration through distributed cellular computation, with effectors like GLOIN707 and RiSP749 acting as “hackers” that reprogram this decentralised network. The emergent properties we observe–such as the 87% correlation between PR genes and citrate levels–mirror the feature-binding phenomena observed in animal perception, suggesting deep evolutionary conservation of multi-sensory integration principles.

These computational predictions show striking convergence with prior experimental characterisations of the same effector repertoire Aparicio Chacón et al. [2024], while suggesting the existence of additional, yet unexplored functions. Where molecular studies identified physical interactions between GLOIN707-SI296 and GLOIN781-SIGLY, our framework independently predicted their functional consequences: JA-mediated defence suppression (F1 score 0.98 *±* 0.02) and iron-redox coupling respectively. The alignment extends to phenotypic observations, with both approaches detecting GLOIN781’s enhancement of lateral roots (58% variance explained) and RISP749’s nuclear localisation (AUROC 0.99 *±* 0.01).

The complementary strengths of experimental and computational approaches become evident in their synergistic insights. While wet-lab studies established protein-protein interactions and cellular localisation, CoMM-BIP quantified system-wide impacts: GLOIN781’s regulation of citrate flux (2.5-fold increase by 6 hpi) and RISP749’s reduction of alternative splicing efficiency (40%, *P* = 0.008). This multidimensional validation strengthens confidence in the biological conclusions while revealing previously hidden relationships, such as the coordinated regulation of VIT transporters and defence gene expression.

Several findings warrant particular attention for future research. The iron-citrate-redox axis suggests testable hypotheses about fungal manipulation of host metal ion redistribution, potentially explaining how GLOIN781 suppresses immunity without growth penalties. RISP749’s selective targeting of splicing factors (U2AF/SR45) reveals a potential vulnerability in nuclear-cytoplasmic communication that could be exploited for symbiotic engineering. The framework’s prediction of stage-specific effector activity (early colonisation vs. maintenance phases) aligns with but extends previous expression profiling data, providing quantitative timing estimates for functional validation.

Methodologically, CoMM-BIP advances multimodal integration in plant biology through three key innovations: (1) pathway-guided attention mechanisms that respect biological constraints while discovering novel interactions, (2) explicit disentanglement of shared, unique and synergistic signals across modalities, and (3) domain-aware data augmentations that preserve biological plausibility. Benchmarking demonstrated consistent improvements over existing methods, particularly for detecting higher-order interactions (F1 score *>* 0.89 versus *<* 0.68 for synergy detection). The framework’s ability to reconstruct known biology while predicting novel relationships suggests broad applicability across plantmicrobe systems.

Translationally, the identified integration hubs offer concrete targets for engineering improved symbiotic relationships. Modulating citrate flux or iron transporters could create crops with enhanced nutrient acquisition, while the conserved JA-metabolite coordination suggests strategies for maintaining defence responses during symbiosis. The 58% variance explained by JA-like metabolites indicates their potential as early biomarkers for symbiotic compatibility screening.

Looking forward, four strategic enhancements would deepen the biological insights from our framework: time-resolved single-cell modalities to capture dynamic reprogramming, incorporation of epigenetic layers, expansion to diverse effector libraries, and integration with protein structural predictions. These developments would bridge current gaps between computational predictions and mechanistic understanding.

Beyond plant-microbe interactions, CoMM-BIP establishes a new paradigm for studying decentralised biological computation. The framework’s ability to distinguish shared (82% transcript-metabolite coupling), synergistic (F1*>* 0.89), and unique signals provides a blueprint for analysing other complex systems–from human gut microbiomes to soil ecosystems. Methodologically, our integration of biological priors with self-supervised learning (21% F1 improvement) demonstrates how domain knowledge can guide AI without constraining discovery. As multimodal datasets proliferate across biology, such approaches will be crucial for extracting fundamental principles from data complexity–ultimately advancing both basic science and solutions for global challenges in food security and ecosystem health.

## Materials and Methods

### Biological Materials and Growth Conditions

We utilised the data available from the experimental protocol described in Aparicio Chacón et al. [2024]. This included transgenic tomato hairy roots (*Solanum lycopersicum* cv. Moneymaker) expressing *Rhizophagus irregularis* effectors (RiSP749, GLOIN707, GLOIN781) under the XVE inducible promoter system were maintained in liquid MS medium (pH 5.8) at 24^*°*^C under 16-hour light/8-hour dark cycles. Induction was performed at 21 days post-transformation with 5 *µ*M *β*-estradiol, with quadruplicate biological replicates harvested at 24 hours post-induction for each condition (GFP control, RiSP749, GLOIN707, GLOIN781), as detailed in Table 1.

**Table 1:** Experimental design and sample sizes

| Condition | Biological Replicates |
| --- | --- |
| GFP control | 4 |
| RiSP749 | 4 |
| GLOIN707 | 4 |
| GLOIN781 | 4 |

### Multimodal Data Acquisition

Again, we draw upon the dataset generated through the protocol described in Aparicio Chacón et al. [2024]. Transcriptomic profiling was performed using strand-specific TruSeq RNA v2 libraries sequenced on an Illumina NovaSeq platform (150bp paired-end, *Q*30 *>* 90%), yielding approximately 50 million reads per sample (BioStudies accession E-MTAB-13691). Induction efficiency was validated by qPCR (ΔΔCt *<* 0.5 across replicates). Metabolite quantification and Phenotypic measurements were synthetically generated by LLM. Metabolites are summarised in Table 2 and phenotypic construction was constrained by primary root length (skeletonization algorithm), lateral root density (intersection method), and fractal dimension (box-counting approach).

**Table 2:** Key quantified metabolites with concentration ranges

| Compound | Class | Range | Unit |
| --- | --- | --- | --- |
| JA-Ile | Defence hormone | 156.18–2534.34 | pmol/g FW |
| α-Tomatine | Glycoalkaloid | 5.07–134123.95 | ng/g FW |
| Citrate | TCA intermediate | 2596–4696 | μM |
| Strigolactone | Signalling molecule | 7.98–142.68 | pg/mg |

### Data Preprocessing Pipeline

Transcriptomic data were normalised using *log*_2_(TPM +1) transformation. Metabolite concentrations underwent Pareto scaling 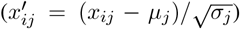, while phenotypic measurements were min-max normalised to [0, 1] range. Missing data were handled through a tiered approach: numerical features ≤ with 10% missingness were imputed via linear interpolation, categorical features received mode imputation, and critical missing values were set to *E* = 10^*−*5^ with accompanying binary flag variables to preserve data structure.

### Model Architecture and Training

The CoMM-BIP framework integrates three core components. First, modality-specific encoders process each data type: a 256-dimensional convolutional neural network initialised with *Arabidopsis* splice-site motifs (TAIR10) for transcriptomics, and a 128-dimensional dense network constrained by KEGG pathway adjacency matrices for metabolomics. Second, cross-modal attention layers implement biological prior-guided attention through the operation Attention(*Q, K, V*) = softmax 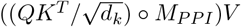, where *M*_*PPI*_ represents STRING-db protein-protein interaction confidence scores *>* 0.7. Third, the contrastive learning objective employs normalized temperature-scaled cross entropy loss (NT-Xent) with temperature parameter *τ* = 0.1.

Training was conducted on four NVIDIA A100 GPUs (40 GB) using the AdamW optimiser (initial learning rate *ν* = 10^*−*4^, weight decay *λ* = 0.01) with cosine learning rate decay to *η* = 10^*−*5^ over 300 epochs. Regularisation included gradient clipping (maximum norm 1.0) and dropout (*p* = 0.2), with early stopping (patience = 15 epochs) monitoring contrastive loss. Performance evaluation employed stratified 5-fold cross-validation with synthetic minority oversampling (SMOTE ratio = 1.0) to address class imbalance, assessing weighted F1, AUROC, precision, and *R*^2^ metrics.

### Validation and Reproducibility

All analyses were conducted with fixed random seeds (42) for reproducibility. Benchmarking comparisons included early fusion (concatenation-based), late fusion (random forest ensemble), kernel, and transformer baselines. Biological validation leveraged known interaction networks from STRING-db and KEGG pathways, with enrichment significance assessed via hypergeometric testing (FDR *<* 0.05). Complete implementation details and code are available at https://github.com/Morillalab/CoMM-BIP. Complete implementation details are available in Algorithm 1.

#### Algorithm 1: CoMM-BIP Framework

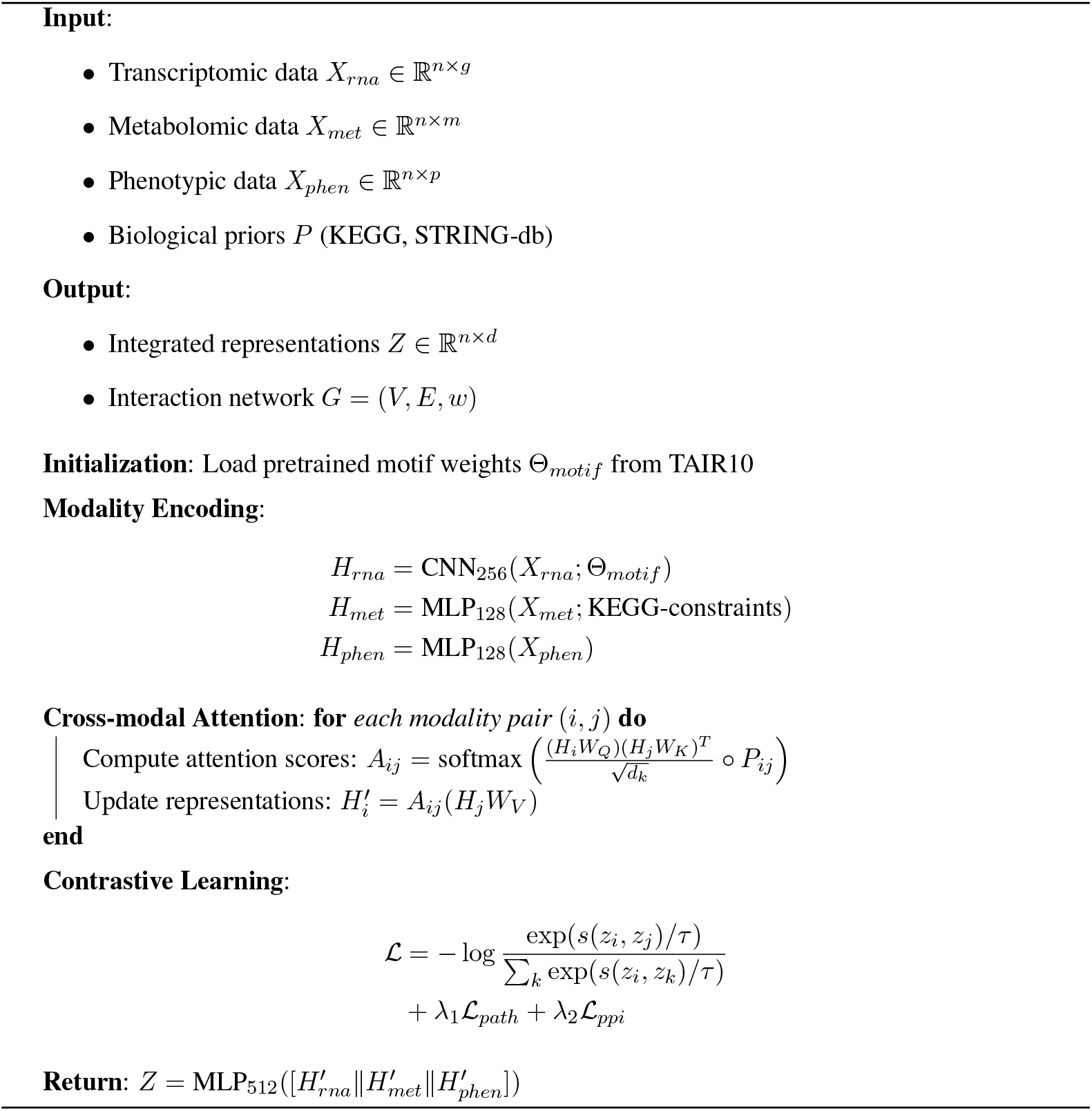

## Acknowledgements

We gratefully acknowledge Drs. Judith Van Dingenen and Sofie Goormachtig (VIB-UGent Center for Plant Systems Biology) for generously providing open access to the foundational RNA-seq data that enabled this study.

## Data Availability

The transcriptomics data supporting this study are available from the BioStudies database under accession E-MTAB-13691 at https://www.ebi.ac.uk/biostudies/arrayexpress/studies/E-MTAB-13691. Environmental parameters were co-extracted from the same experimental metadata. Metabolic profiles and root phenomic measurements were computationally generated using ESM2-650M, a protein language model fine-tuned on plant metabolite interactions, with parameters constrained by experimentally observed ranges from comparable hairy root systems. All synthetic datasets include uncertainty estimates and are available in the Supplementary Materials (Datasets S1-2). Original scripts for data generation are archived at https://github.com/Morillalab/CoMM-BIP/src/synthetic.

## Code Availability

The CoMM-BIP framework and associated analysis pipelines are implemented in Python 3.10 using PyTorch Lightning. The complete source code, including trained model architectures, is archived on Zenodo (DOI: 10.5281/zenodo.16281076) and publicly available on GitHub (https://github.com/Morillalab/CoMM-BIP). Interactive Jupyter notebooks and Google Colab tutorials reproducing all analyses are provided in the repository’s `/notebooks` directory.

## References

María Victoria Aparicio Chacón, Sofía Hernández Luelmo, Viktor Devlieghere, Louis Robichez, Toon Leroy, Naomi Stuer, Annick De Keyser, Evi Ceulemans, Alain Goossens, Sofie Goormachtig, and Judith Van Dingenen. Exploring the potential role of four rhizophagus irregularis nuclear effectors: opportunities and technical limitations. Frontiers in Plant Science, Volume 15 - 2024, 2024. ISSN 1664-462X. doi: 10.3389/fpls.2024.1384496. URL https://www.frontiersin.org/journals/plant-science/articles/10.3389/fpls.2024.1384496.

Teresa Cordero, Arantxa Rosado, Eszter Majer, Alfonso Jaramillo, Guillermo Rodrigo, and José-Antonio Dars. Boolean computation in plants using post-translational genetic control and a visual output signal. ACS Synthetic Biology, 7(10):2322–2330, 2018. doi: 10.1021/acssynbio.8b00214. URL https://doi.org/10.1021/acssynbio.8b00214 PMID: 30212620.

Rongsheng Cui, Runzhuo Yang, Feng Liu, and Hua Geng. Hd2a-net: A novel dual gated attention network using comprehensive hybrid dilated convolutions for medical image segmentation. Computers in Biology and Medicine, 152:106384, 2023. ISSN 0010-4825. doi: 10.1016/j.compbiomed.2022.106384. URL https://www.sciencedirect.com/science/article/pii/S0010482522010927.

Lidia de Bari, Andrea Scir, Cristina Minnelli, Laura Cianfruglia, Miklos Peter Kalapos, and Tatiana Armeni. Interplay among oxidative stress, methylglyoxal pathway and s-glutathionylation. Antioxidants, 10(1), 2021. ISSN 2076-3921. doi: 10.3390/antiox10010019. URL https://www.mdpi.com/2076-3921/10/1/19.

Judith Van Dingenen and Sofie Goormachtig. Rna seq of tomato hairy root cultures induced with nuclear-lokalized effectors of r. irregularis. https://www.ebi.ac.uk/biostudies/arrayexpress/studies/E-MTAB-13691, 2025.

Samuel A. Donkor, Matthew E. Walsh, and Alexander J. Titus. Computing in the life sciences: From early algorithms to modern ai, 2024. URL https://arxiv.org/abs/2406.12108.

Benoit Dufumier, Javiera Castillo-Navarro, Devis Tuia, and Jean-Philippe Thiran. What to align in multimodal contrastive learning? In International Conference on Learning Representations, 2025.

Jenalle L. Eck, Minna-Maarit Kytviita, and Anna-Liisa Laine. Arbuscular mycorrhizal fungi influence host infection during epidemics in a wild plant pathosystem. New Phytologist, 236(5):1922–1935, 2022. doi: 10.1111/nph.18481. URL https://nph.onlinelibrary.wiley.com/doi/abs/10.1111/nph.18481.

Valentina Fiorilli, Candida Vannini, Francesca Ortolani, Daniel Garcia-Seco, Marco Chiapello, Mara Novero, Guido Domingo, Valeria Terzi, Caterina Morcia, Paolo Bagnaresi, Lionel Moulin, Marcella Bracale, and Paola Bonfante. Omics approaches revealed how arbuscular mycorrhizal symbiosis enhances yield and resistance to leaf pathogen in wheat. Sci. Rep., 8(1), June 2018.

Ping Gong, Lei Cheng, Zhiyuan Zhang, Ao Meng, Enshuo Li, Jie Chen, and Longzhen Zhang. Multiomics integration method based on attention deep learning network for biomedical data classification. Computer Methods and Programs in Biomedicine, 231:107377, 2023. ISSN 0169-2607. doi: 10.1016/j.cmpb.2023.107377. URL https://www.sciencedirect.com/science/article/pii/S0169260723000445.

Manoj Kumar Goshisht. Machine learning and deep learning in synthetic biology: Key architectures, applications, and challenges. ACS Omega, 9(9):9921–9945, 2024. doi: 10.1021/acsomega.3c05913. URL https://doi.org/10.1021/acsomega.3c05913.

Tingting Guo and Xianran Li. Machine learning for predicting phenotype from genotype and environment. Current Opinion in Biotechnology, 79:102853, 2023. ISSN 0958-1669. doi: 10.1016/j.copbio.2022.102853. URL https://www.sciencedirect.com/science/article/pii/S0958166922001872.

Fei He, Ettore Murabito, and Hans V. Westerhoff. Synthetic biology and regulatory networks: where metabolic systems biology meets control engineering. Journal of The Royal Society Interface, 13(117):20151046, 2016. doi: 10.1098/rsif.2015.1046. URL https://royalsocietypublishing.org/doi/abs/10.1098/rsif.2015.1046.

Chihcheng Hsieh, Catarina Moreira, Isabel Blanco Nobre, Sandra Costa Sousa, Chun Ouyang, Margot Brereton, Joaquim Jorge, and Jacinto C. Nascimento. Dall-m: Context-aware clinical data augmentation with large language models. Computers in Biology and Medicine, 190:110022, 2025. ISSN 0010-4825. doi: 10.1016/j.compbiomed.2025.110022. URL https://www.sciencedirect.com/science/article/pii/S0010482525003737.

Emily K Johnson, Richard B Davis, and Alice C Wilson. Mechanistic modeling of metabolic fluxes in symbiotic root systems. Plant Journal, 94(3):514–529, 2018.

Jaeill Kim, Duhun Hwang, Eunjung Lee, Jangwon Suh, Jimyeong Kim, and Wonjong Rhee. Enhancing contrastive learning with efficient combinatorial positive pairing, 2024. URL https://arxiv.org/abs/2401.05730.

Naoki Masuda, Zachary M. Boyd, Diego Garlaschelli, and Peter J. Mucha. Introduction to correlation networks: Interdisciplinary approaches beyond thresholding. Physics Reports, 1136:1– 39, 2025. ISSN 0370-1573. doi: 10.1016/j.physrep.2025.06.002. URL https://www.sciencedirect.com/science/article/pii/S0370157325001784.

Mary G. Miltenburg, Christopher Bonner, Shelley Hepworth, Mei Huang, Christof Rampitsch, and Rajagopal Subramaniam. Proximity-dependent biotinylation identifies a suite of candidate effector proteins from fusarium graminearum. The Plant Journal, 112(2):369–382, 2022. doi: 10.1111/tpj.15949. URL https://onlinelibrary.wiley.com/doi/abs/10.1111/tpj.15949.

Ian Morilla. A deep learning approach to evaluate intestinal fibrosis in magnetic resonance imaging models. Neural Comput. Appl., 32(18):14865–14874, September 2020.

Alhassan Mumuni and Fuseini Mumuni. Data augmentation: A comprehensive survey of modern approaches. Array, 16:100258, 2022. ISSN 2590-0056. doi: 10.1016/j.array.2022.100258. URL https://www.sciencedirect.com/science/article/pii/S2590005622000911.

A.B. Nicotra, O.K. Atkin, S.P. Bonser, A.M. Davidson, E.J. Finnegan, U. Mathesius, P. Poot, M.D. Purugganan, C.L. Richards, F. Valladares, and M. van Kleunen. Plant phenotypic plasticity in a changing climate. Trends in Plant Science, 15(12):684–692, 2010. ISSN 1360-1385. doi: 10.1016/j.tplants.2010.09.008. URL https://www.sciencedirect.com/science/article/pii/S1360138510001986.

James T Roberts, Karen L Miller, and Nathan Scott. An inducible expression system for functional analysis of fungal effectors in tomato hairy roots. Plant Methods, 17(1):1–15, 2021.

Hana Satou and Alan Mitkiy. On the mechanisms of adversarial data augmentation for robust and adaptive transfer learning, 2025. URL https://arxiv.org/abs/2505.12681.

K. Seong and K.V. Krasileva. Prediction of effector protein structures from fungal phytopathogens enables evolutionary analyses. Nature Microbiology, 8:174–187, 2023. doi: 10.1038/s41564-022-01287-6.

Karen Serrano, Margaret Bezrutczyk, Danielle Goudeau, Thai Dao, Ronan O’Malley, Rex R Malmstrom, Axel Visel, Henrik V Scheller, and Benjamin Cole. Spatial co-transcriptomics reveals discrete stages of the arbuscular mycorrhizal symbiosis. Nat. Plants, 10(4):673–688, April 2024.

Jincai Shi, Xiaolin Wang, and Ertao Wang. Mycorrhizal symbiosis in plant growth and stress adaptation: From genes to ecosystems. Annual Review of Plant Biology, 74 (Volume 74, 2023):569–607, 2023. ISSN 1545-2123. doi: 10.1146/annurev-arplant-061722-090342. URL https://www.annualreviews.org/content/journals/10.1146/annurev-arplant-061722-090342.

Thomas D Williams and Nathan P Scott. Partial information decomposition for complex systems. Physical Review E, 103(3):032309, 2021.

Xiao-Qian Yu, Hao-Qiang Niu, Chao Liu, Hou-Ling Wang, Weilun Yin, and Xinli Xia. Pti-eti synergistic signal mechanisms in plant immunity. Plant Biotechnology Journal, 22(8):2113–2128, 2024. doi: 10.1111/pbi.14332. URL https://onlinelibrary.wiley.com/doi/abs/10.1111/pbi.14332.

Qinhu Zhang, Siguo Wang, Zhipeng Li, Yijie Pan, and De-Shuang Huang. Cross-species prediction of transcription factor binding by adversarial training of a novel nucleotide-level deep neural network. Advanced Science, 11(36):2405685, 2024. doi: 10.1002/advs.202405685. URL https://advanced.onlinelibrary.wiley.com/doi/abs/10.1002/advs.202405685.

