## Supplementary figures and images for "Multimodal Learning Reveals Plants’ Hidden Sensory Integration Logic"

### Figure S1

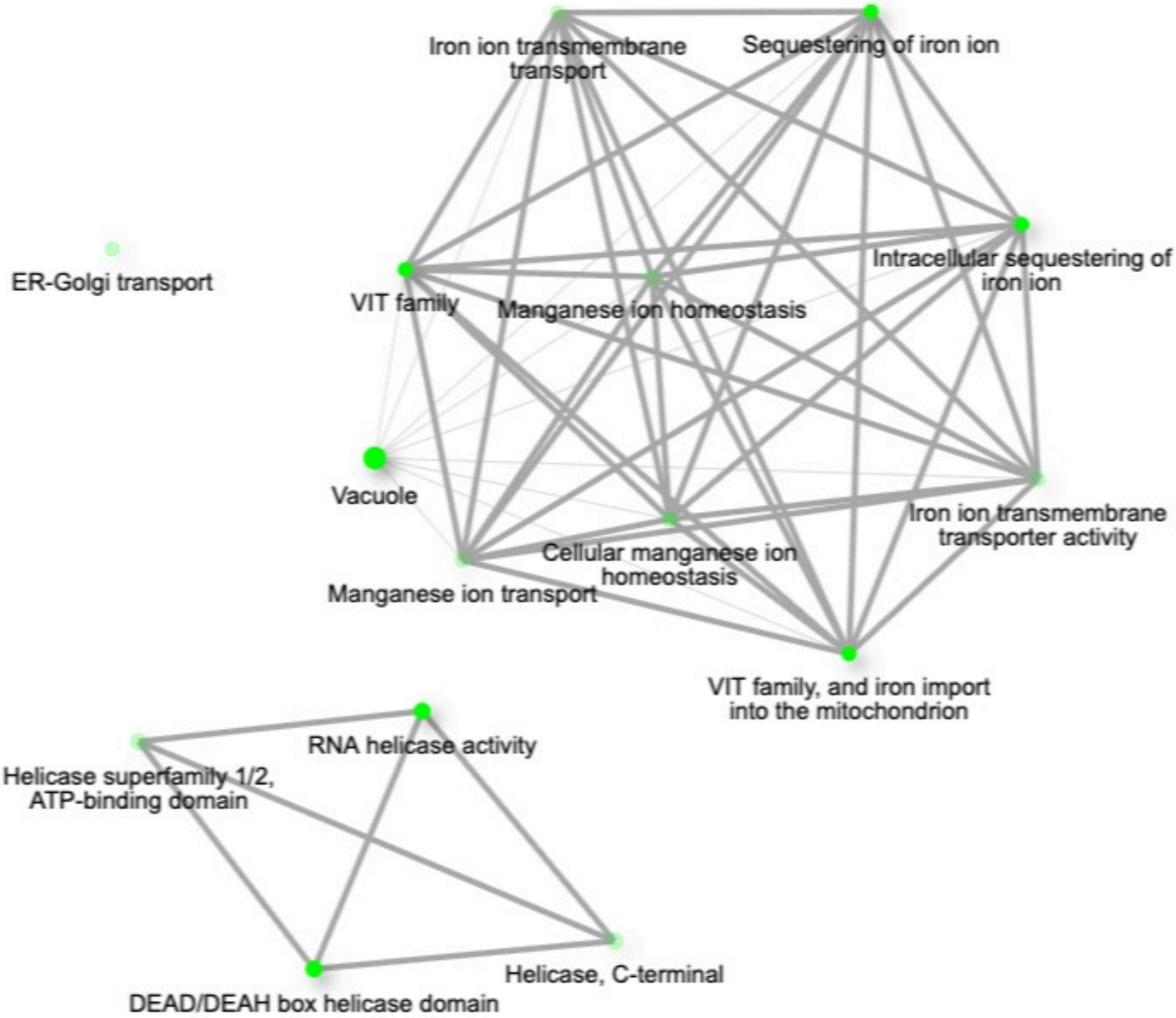

### Figure S2

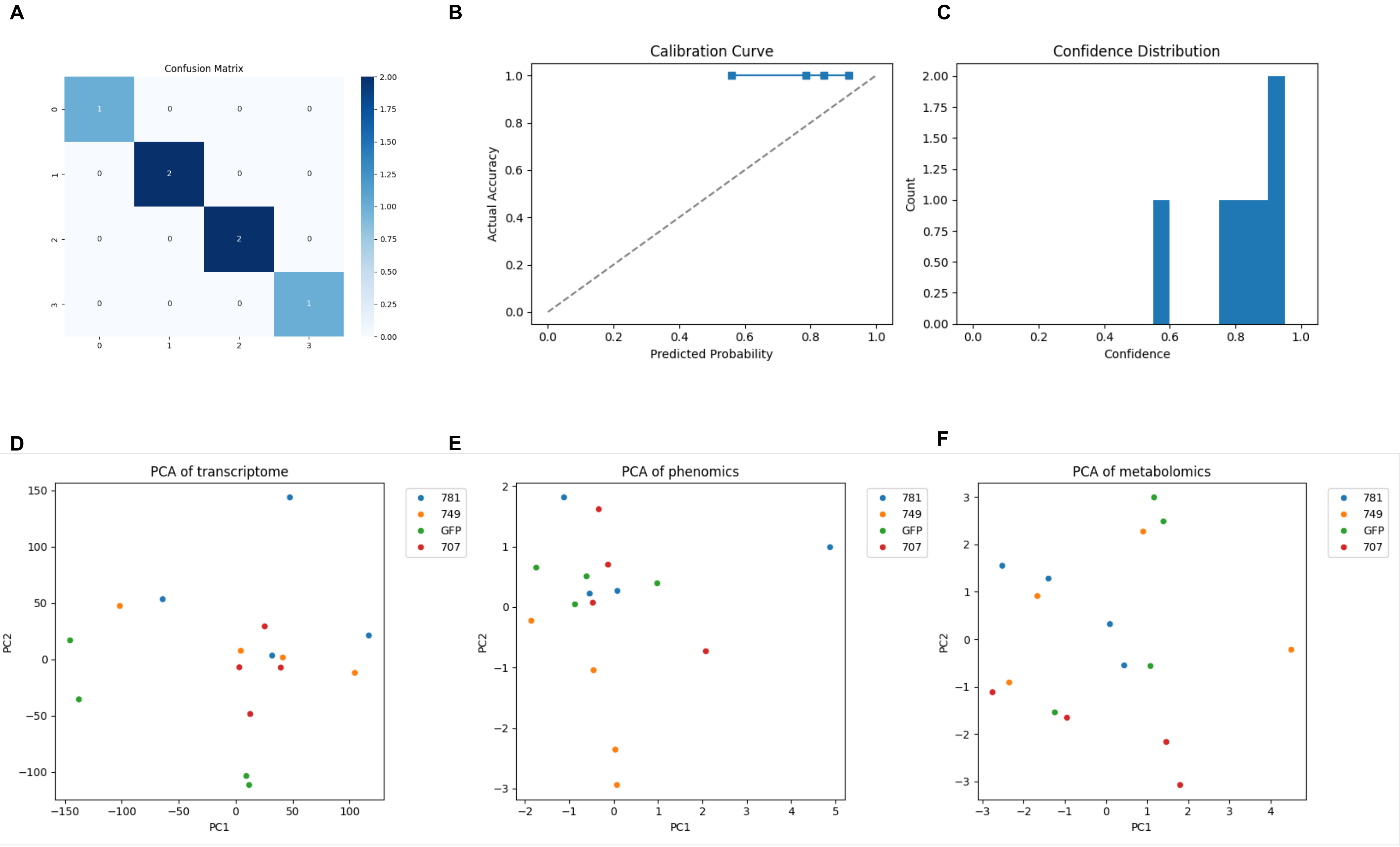
