## Supplementary material for "Multimodal Learning Reveals Plants’ Hidden Sensory Integration Logic": Figure S3

**A. Phenotypic Regression ($r^2$)**

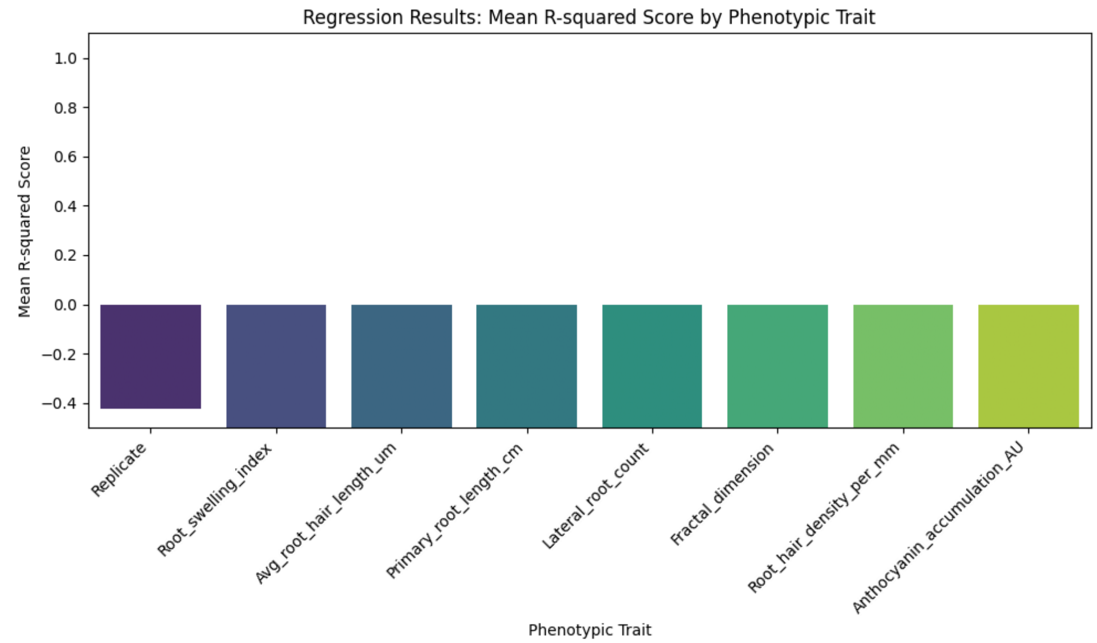


**B. Phenotypic Regression (*MSE*)**

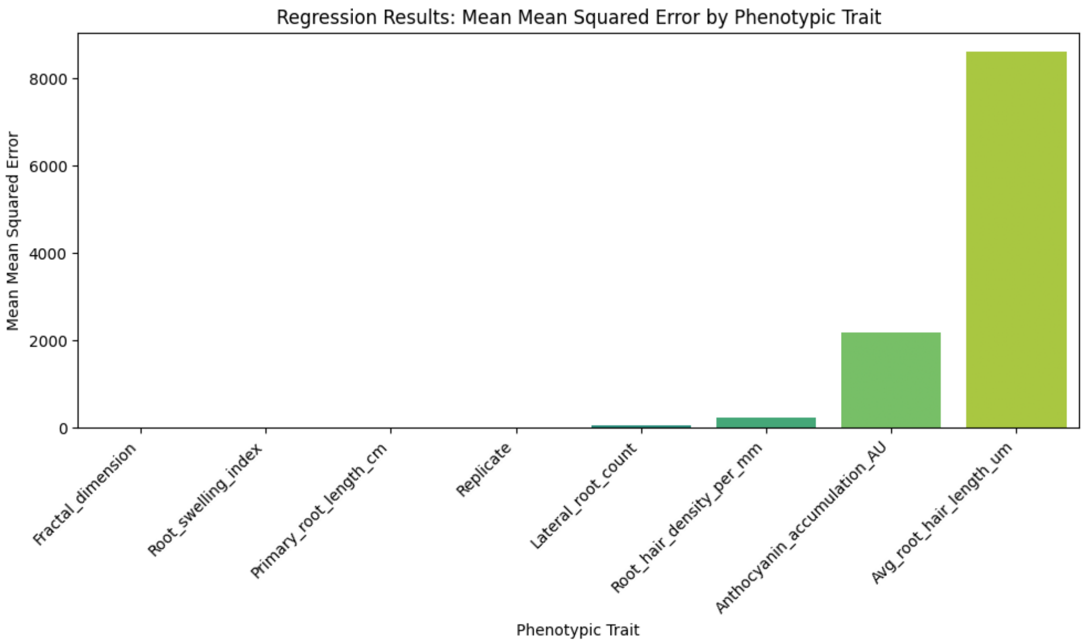


**C. Genotype Classification by Embeddings**

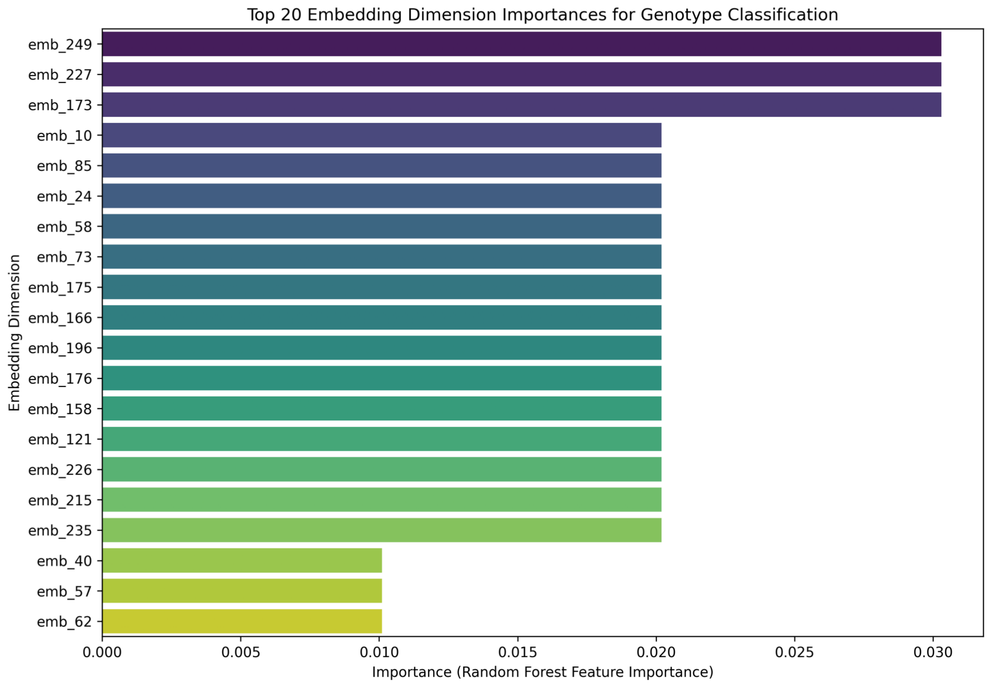


**D. Embeddings Interpretability by Effector**

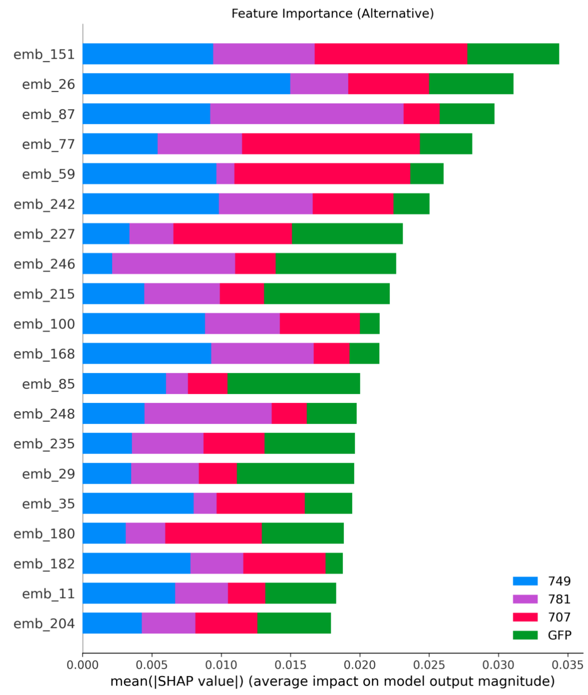
